# Cytosolic delivery of anionic cyclic dinucleotide STING agonists with locally supercharged viral capsids

**DOI:** 10.1101/2024.10.07.617100

**Authors:** Paul Huang, Paige E. Pistono, Hannah S. Martin, Jennifer L. Fetzer, Matthew B. Francis

## Abstract

Cyclic dinucleotide (CDN) STING agonists represent a powerful new immunotherapy treatment modality and are a class of nucleotide-based therapies with broad clinical potential. However, they face practical challenges for administration, largely due to their poor pharmacological properties. We report the development of a drug delivery platform for CDNs and other anionic small-molecule drugs using bacteriophage MS2 viral capsids with engineered cationic residues. Relative to viral capsids lacking locally supercharged regions, these assemblies exhibit substantial increases in mammalian cell uptake while avoiding cell toxicity and hemolysis. CDN drugs were covalently attached to the interior capsid surfaces through reductively cleavable disulfide linkers, which allowed for traceless drug release upon cell entry and exposure to reductive cytosolic environments. MS2-mediated CDN delivery into immune cell populations resulted in an approximately 100-fold increase in delivery efficiency compared to free drugs and showed enhanced STING activation as well as downstream cytokine release.

## INTRODUCTION

Nucleotides and other negatively charged drugs are highly promising therapeutic modalities with a wide range of potential applications, including in gene therapy, immune modulation, vaccination, and genetic activation and inhibition^1^. However, the clinical use of such approaches for precise intracellular targets has been largely precluded due to the poor pharmacological behavior of anionic nucleotide-based drugs. These compounds stray beyond conventional “drug-like” properties and therefore suffer from low bioavailability due to impermeability across the cell membrane, degradation by nucleases, and rapid clearance from the bloodstream^2,3^. As the anionic character of such drugs is often essential for tight substrate binding and efficacy, these challenges have been difficult to overcome using traditional medicinal chemistry approaches. As a result, a number of labs are developing nanoscale drug delivery platforms to improve the bioavailability and consequently the efficacy of nucleotide-based anionic drugs^3–5^.

One particular focus in anionic nucleotide therapies involves the cGAS-STING pathway, an innate immune signaling pathway that senses cytosolic DNA as a result of cell damage or viral pathogen invasion. This pathway is of special interest in anticancer immunotherapy due to its potential to activate and target the immune system against tumor cells^6–8^. The cGAS-STING pathway has been implicated in several tumor clearance mechanisms, including in radiation therapy^9^ and DNA-targeted chemotherapy^10^. Specific pharmaceutical efforts have focused on mimicking the small molecule cyclic dinucleotide (CDN) 2’3’-cyclic GMP-AMP, or cGAMP^11^, which is synthesized by the DNA-sensing protein cGAS and functions as a second messenger to bind and activate the STING pathway^12–14^. This activation step, however, requires CDNs to bind to the STING receptor located on the cytosolic face of the endoplasmic reticulum (ER). Because anionic CDNs cannot freely cross the cell membrane, they rely on an assortment of relatively lower-affinity membrane transporters that are variably expressed across cell types and can be downregulated in cancers^15–18^. As a result, while many synthetic CDN analogs have been developed with STING agonist activity, to date all have faced challenges in clinical trials in large part due to poor cellular uptake, premature degradation, and/or toxicity at high doses^19,20^. While some more membrane-permeable non-anionic STING agonists have been developed^21^, their mechanisms of action are yet not well understood, especially regarding how they differ from natural CDNs. Among nanoscale drug delivery vehicles, work has been done to adapt oligonucleotide carriers, such as liposomes^22,23^, polymer-based nanoparticles^24^, hydrogels^25^, and extracellular vesicles^26^, for use in cyclic dinucleotide drug delivery.

Virus-like particles (VLPs) are naturally well-suited for nucleotide drug delivery, and previous mechanistic studies of host cell immune responses to viral infections have observed the ability of some enveloped native viruses and VLPs to package cGAMP and transfer it between cells^27,28^. This strategy has been demonstrated for the delivery of cGAMP to tumor cells both *in vitro*^*28*^ and *in vivo*^*29*^. However, the delivery of STING agonists using synthetic VLPs, which are non-infectious but retain the advantageous size and sequestration properties of native viruses, has yet to be reported. In this work, we have adapted bacteriophage MS2, a protein-based non-enveloped VLP, as a covalent carrier for cyclic dinucleotide drugs. This system offers distinct advantages due to its well-mapped structural mutability^30^, as well as its ability to encapsulate and deliver drug cargo of interest to cells. Moreover, we have recently shown that cationic residues can be placed in specific locations on the outer surface of MS2 VLPs to enhance cellular uptake and cytosolic delivery of drug cargo^31,32^. Polycationic drug delivery agents have long been explored as cell-penetrating agents, due to their association with anionic glycan residues on cell surfaces and disruption of endosomal membranes^33–36^. However, their globally “supercharged” properties (and thus net positive charges) often leave them susceptible to systemic toxic effects^37^. Guided by deep mutational scanning studies, careful engineering of positive charges on the FG loop of MS2 produced localized patches of concentrated positive charge from the introduction of just two cationic residue mutations per monomer. This conservative change was found to enhance cellular uptake by up to 67-fold with minimal alterations to the global negative charge of the capsid. In contrast to many fully cationic carriers, these capsids showed minimal cell toxicity or hemolysis even at high exposure concentrations.

Here, we report that these locally supercharged capsids not only improve uptake by the cell but also facilitate the transfer of membrane-impermeable cyclic dinucleotides into the cell to access the cytosolic-facing STING receptor. The drug molecules are attached inside the capsid by virtue of a covalent linkage and liberated in the cells through a glutathione-based release platform. This system was able to achieve the effective delivery of CDN drugs to cells, resulting in up to a 100-fold EC_50_ improvement in downstream STING activation over the free drugs. These results demonstrate that locally supercharged MS2 capsids can successfully achieve intracellular delivery without compromising the native affinity of CDNs to the STING protein, thus providing a promising new platform that could expand the therapeutic window of this important class of STING agonist drugs.

## RESULTS

### Locally charged MS2 capsids enhance delivery of anionic cargo

The capsid structure of native bacteriophage MS2 consists of 180 identical coat proteins, arranged in a hollow 27 nm sphere with 32 pores that are 2 nm in size^38^. This latter feature allows for passive diffusion of small molecules in and out of the capsid interior but restricts passage of proteins and other macromolecules. We have previously engineered a non-native cysteine residue into a mutable site at position 87 in the capsid interior, which we have used to attach small-molecule drug cargo such as paclitaxel^39^, doxorubicin^40^, MMAE^31^, and photodynamic therapy agents^41^. In developing locally supercharged MS2 variants, we used our understanding of MS2 capsid structure and the influence of site-specific mutations on overall structural stability to identify compatible sites for cationic residue mutations. The two best-performing variants involved lysine and arginine mutations at exterior-facing positions 71 and 73, which cluster at the 5- and 6-fold assembly interfaces (**Figure 1A**). This creates localized pockets of positive charge on the exterior surface of each capsid. These two mutants, bearing T71K/G73R and T71R/G73K mutations, exhibited enhanced cellular uptake of a covalently attached fluorescein maleimide dye. These uptake properties can also be observed by treating cultured cells with MS2 attached to an anionic AlexaFluor 594 maleimide dye, which does not enter or associate with the cell as a free small molecule in solution (**Figure 1B**). These results also demonstrate that covalent interior attachment of anionic cargo does not adversely affect the delivery properties of these MS2 capsids. While the precise mechanism of cell uptake is still being studied, it is believed that the cell uptake process is initiated from the higher affinity of the capsids to anionic cell-surface residues, such as glycosaminoglycans^31^.

**Figure 1.**
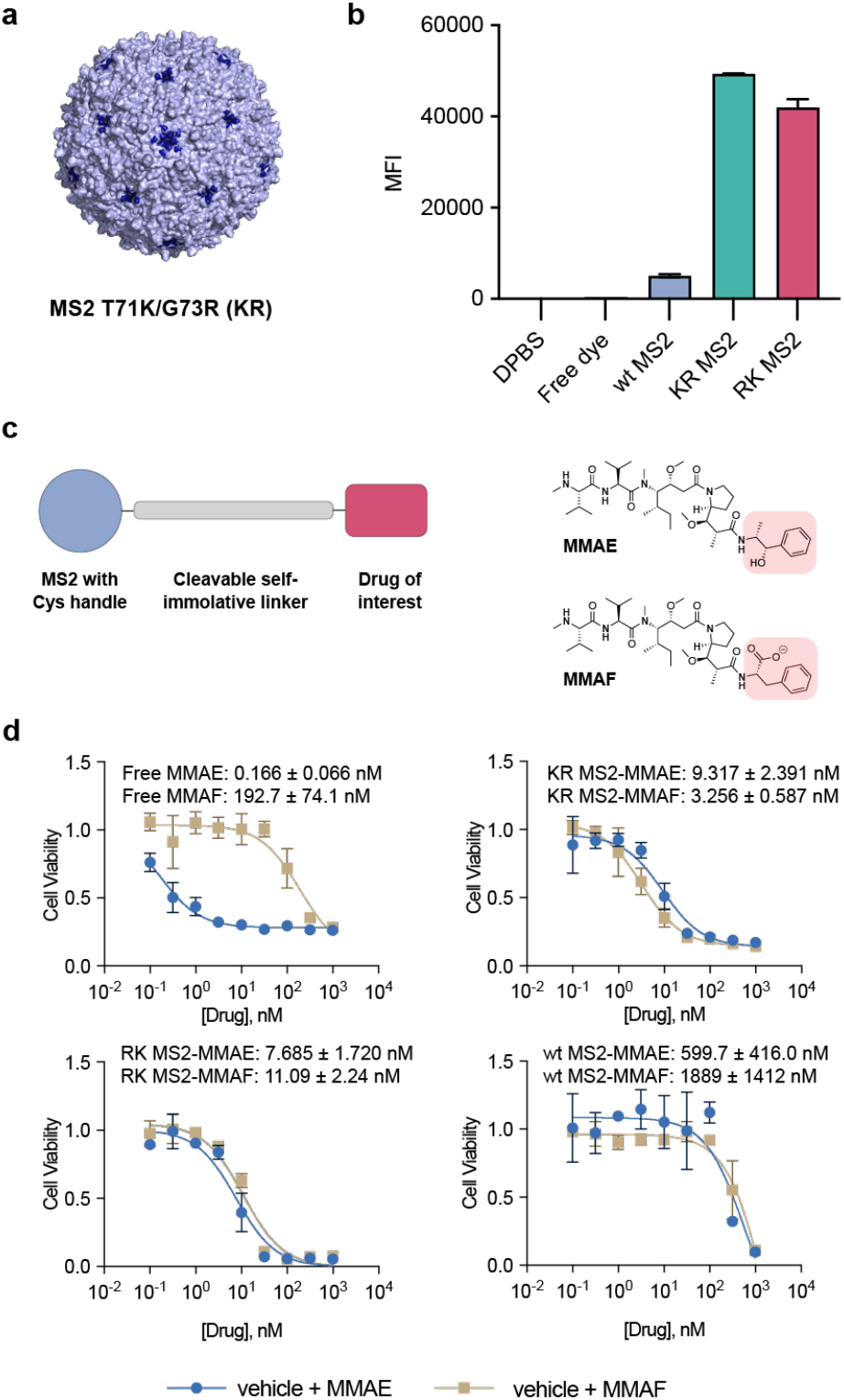
Locally supercharged MS2 capsids for intracellular delivery. **(a)** Structure of bacteriophage MS2 coat protein (PDB: 2MS2) with mutations T71K and G73R shown in blue. **(b)** Cellular uptake of 5 µM MS2-conjugated or free AlexaFluor 594 into U-251-MG cells. **(c)** MS2-drug conjugate scheme, and chemical structures of MMAE and MMAF with structural differences emphasized. **(d)** MTS cell viability assay of U-251-MG cells treated with a range of doses of MMAE and MMAF, either free in solution or conjugated to MS2 through a cleavable valine-citrulline linker. Cells were treated with free or MS2-delivered drug for 72 h before assessing viability. The resulting survival curves are shown with EC_50_ values.

In a previous report, we used the T71K/G73R (KR MS2) and T71R/G73K (RK MS2) capsids to deliver monomethyl auristatin E (MMAE) to cancer cells, demonstrating enhanced cell killing relative to MMAE-loaded wild-type MS2^31,32^. In the studies we report here, we began by comparing the delivery of MMAE to that of the structural analogue MMAF. Both drugs operate via inhibition of microtubule formation through tight binding of tubulin monomers, but MMAF possesses an anionic charge (**Figure 1C**) that vastly reduces its potency by inhibiting membrane permeability. This has been established both as a free drug and as a payload in antibody drug conjugates that lack endosomal escape mechanisms. To validate the efficacy of the MS2 delivery systems, we treated cancer cell lines with MMAE and MMAF, both as free drugs and as conjugates attached to the interior surfaces of both native and locally supercharged MS2 capsids via an endosomally cleavable valine-citrulline linker. While free MMAF showed considerably reduced potency compared to free MMAE, delivery of both drugs with KR MS2 and RK MS2 showed identically high cytotoxicity (**Figure 1D**). On the other hand, delivery by wild-type MS2 was ineffective except at high drug concentrations, confirming that the positive charge mutations on the capsids are essential for cellular delivery of the anionic MMAF cargo. Taken together, these results demonstrate that these locally supercharged MS2 capsids can both associate with cells to enhance uptake and facilitate the delivery of anionic cargo.

### A fluorescent turn-on dye demonstrates glutathione-triggered cytosolic release of anionic cargo

With the goal of STING agonist delivery in mind, we next designed a cleavable covalent drug delivery platform that could be adapted to cyclic dinucleotides. As MS2 VLPs are porous to small molecules and lack a lipid envelope, loading of small-molecule drug cargo must be done through covalent attachment to the MS2 protein surface. We chose a disulfide release strategy, which takes advantage of the reducing environment of the cell cytoplasm from a high concentration of reduced glutathione^42^. This system was preferable to the valine-citrulline linker due to its release in the cytosol rather than the endosome, as well as being more synthetically straightforward. Furthermore, cargo encapsulation inside MS2 is hypothesized to protect the drugs from disulfide reduction while in transport, where reduction occurs largely through albumin and other proteins too large to pass through the MS2 pores. A self-immolative linker was used to ensure traceless release of drug cargo, ensuring that extraneous covalent groups do not adversely affect the binding of cyclic dinucleotides to STING (**Figure S1A**).

As a proof of concept to test delivery and reductive drug release, we first used a fluorescent platform based on the 4-amino-1,8-naphthalimide dye^43^. While this dye ordinarily fluoresces brightly in the green wavelength region, acylation of the 4-amino position greatly reduces that fluorescence (**Figure 2A, Figure S1B**). Caging of the 4-amino group functions as a proxy to attaching nucleotide drugs via the exocyclic amine of adenosine. We therefore synthesized a napthalimide dye with an added anionic phosphate group and appended the reductively cleavable linker at the 4-amino position (**Figure 2B**) to observe fluorescence turn-on upon glutathione-mediated release. Using an activated 2-thiopyridyl group, the linker-caged dye was attached to the internal cysteine of MS2 with high levels of modification, as monitored by ESI-QTOF mass spectrometry (**Figure 2B**). By monitoring the fluorescence emission, the MS2-dye conjugates were observed to be relatively stable over 10 h at 37 °C. Treatment with 10 mM glutathione in sufficiently pH 7.2-buffered phosphate solution showed a rapid fluorescence increase over several hours (**Figure 2C**), with fluorescence intensity reaching half-maximum at about 2 h. Mass spectrometry analysis of the capsid also showed a similar timeline for detachment of the dye, suggesting that the self-immolative linker cleaved rapidly upon glutathione-mediated liberation from the capsid interior surface (**Figure S1C**).

**Figure 2.**
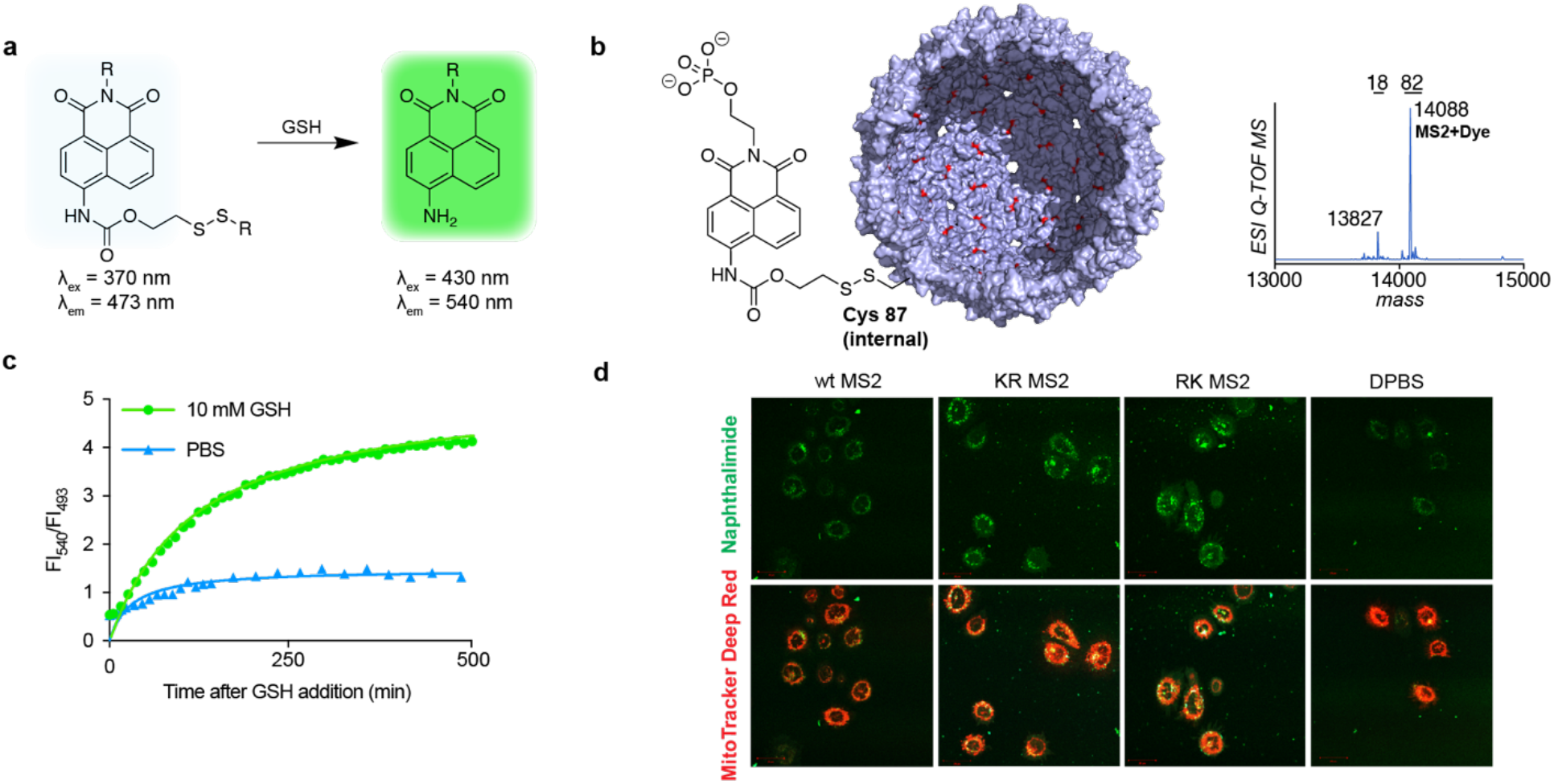
Validation of GSH-mediated release using a naphthalimide reporter dye. **(a)** The 4-amino-1,8-naphthalimide dye fluoresces in the green, but exhibits blue-shifted and significantly reduced fluorescence upon caging of the amino functional group. Addition of glutathione (GSH) liberates the free green dye. **(b)** An anionic phosphate-functionalized dye was caged with a disulfide linker and attached to an engineered cysteine in the interior of MS2. Modification percentage was determined from mass spectra by integration of deconvoluted mass peaks, marked above each peak, with shown 82% dye modification translating to >144 drug copies per capsid. **(c)** A solution of 10 µM dye-conjugated MS2 in PBS was treated with a 10 mM solution of GSH or a PBS control. Fluorescence was monitored over time as a readout for dye release. **(d)** HeLa cells were treated with 5 µM dye-conjugated MS2 bearing or lacking the locally supercharging T71K/G73R (KR) or T71R/G73K (RK) mutations at 37 °C. A 4 h data point is shown. Cells were then stained with MitoTracker™ and imaged live.

We next confirmed that the platform could be used in cells, and that enhanced uptake of the locally supercharged capsids in cancer cell culture could be visualized. Cells treated with both wild-type and locally supercharged MS2 capsids loaded with the linker-caged dye were visualized by confocal fluorescence microscopy. Fluorescence turn-on was observed in the locally supercharged MS2-treated cells, while the native MS2-treated cells exhibited only weak background autofluorescence that was similar to that of untreated cells (**Figure 2D**). This result demonstrates the suitability of the disulfide release platform for delivery and release of anionic small-molecule cargo within the cell.

### Cyclic nucleotides can be adapted to the reductively cleaved linker platform

Following the success of the disulfide linker in cells, we adapted the strategy to a synthetically derived cyclic dinucleotide drug candidate—a phosphorothioate analogue of cyclic di-AMP. The same pyridyl disulfide linker used with the fluorescent turn-on dye was installed on the adenosine base (**Figure 3A**). Of synthetic note, pyridine was found to be the optimal solvent to dissolve the CDN and facilitate the reaction, and modification of both adenosine groups was minimized through short reaction times, with the chromatographic separation of the singly modified product from the unmodified and doubly modified CDN being aided by an ion-pairing buffer. The linker-appended CDN was attached to the MS2 capsid interior via the internal engineered cysteine 87, and ESI-QTOF-MS analysis showed high modification levels around 90% (**Figure 3B-C, Figure S2**). It is worth noting that complete modification of each MS2 monomer would result in a drug loading of 180 drug copies per viral capsid; thus, we estimate that the CDN capsids each possessed approximately 160-168 drug molecules per capsid. Analysis by mass spectrometry and HPLC also showed the re-emergence of the unmodified MS2 capsid as well as the traceless CDN drug after 24 h treatment with 10 mM reduced glutathione (**Figure 3C-D**).

**Figure 3.**
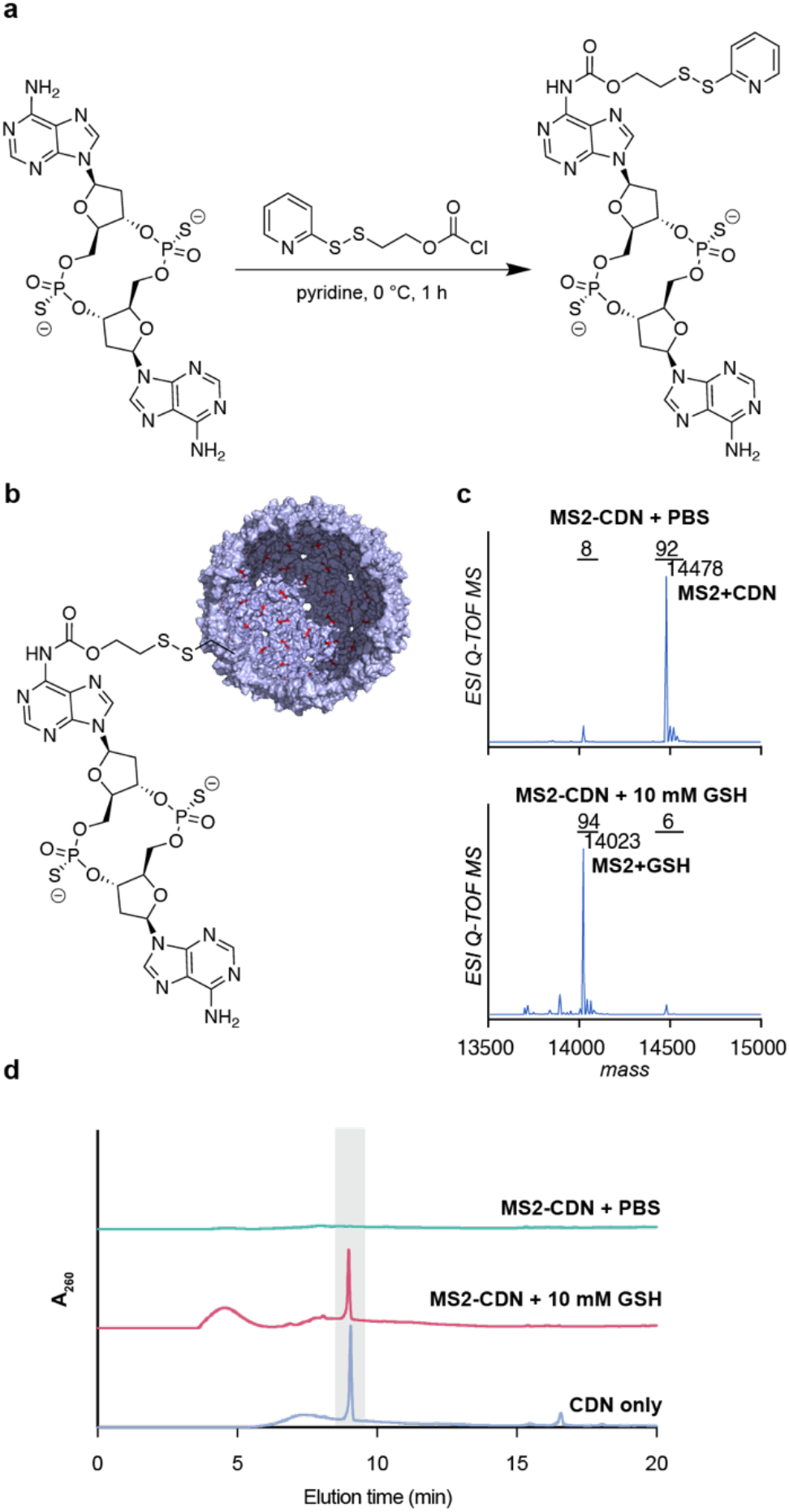
**(a)** Synthetic scheme for disulfide linker attachment to the CDN STING agonist. **(b)** Conjugation via disulfide exchange of the linker-appended CDN to cysteine 87 within the MS2 capsid. **(c)** Protein MS analysis of MS2-CDN conjugate before and after 24 h treatment with 10 mM GSH. **(d)** HPLC analysis of GSH reaction, with the gray box representing the free CDN elution time.

### MS2-delivered CDNs greatly enhance STING activation and downstream immune signaling in cells

Intracellular delivery was next tested using a THP-1 monocyte cell line containing a fluorescent reporter gene that expresses the red fluorescent protein tdTomato upon stimulation of an interferon stimulatory response element (ISRE), which would occur upon STING activation by the CDN drug^16^. These cells were treated with free CDN or MS2-delivered CDN drug for 24 h and analyzed via flow cytometry (**Figure 4A-C**). The free CDN showed an EC_50_ value of ∼1 µM, which compared well to values previously reported in the literature. In contrast, delivery with locally supercharged MS2s achieved an 80-100-fold increase in EC_50_ potency, reaching levels as low as 10-20 nM. No observed STING activation was observed from treatment with drug-free capsids, supporting that the release of the drug, rather than any potential nucleic acid contamination from MS2 expression, was the cause of STING activation (**Figure 4C**). Furthermore, wild-type MS2 capsids loaded with CDN did not meaningfully activate STING in cells, demonstrating the criticality of the cationic mutations in the capsid for cellular uptake and delivery to intracellular STING. Additionally, while some enveloped VLPs can naturally package and encapsulate cyclic dinucleotides^27–29^, treatment of THP-1 cells and CDN mixtures *in trans* yielded no significant improvement over free drug alone, indicating that covalent attachment is required for MS2-mediated delivery (**Figure S3**).

**Figure 4.**
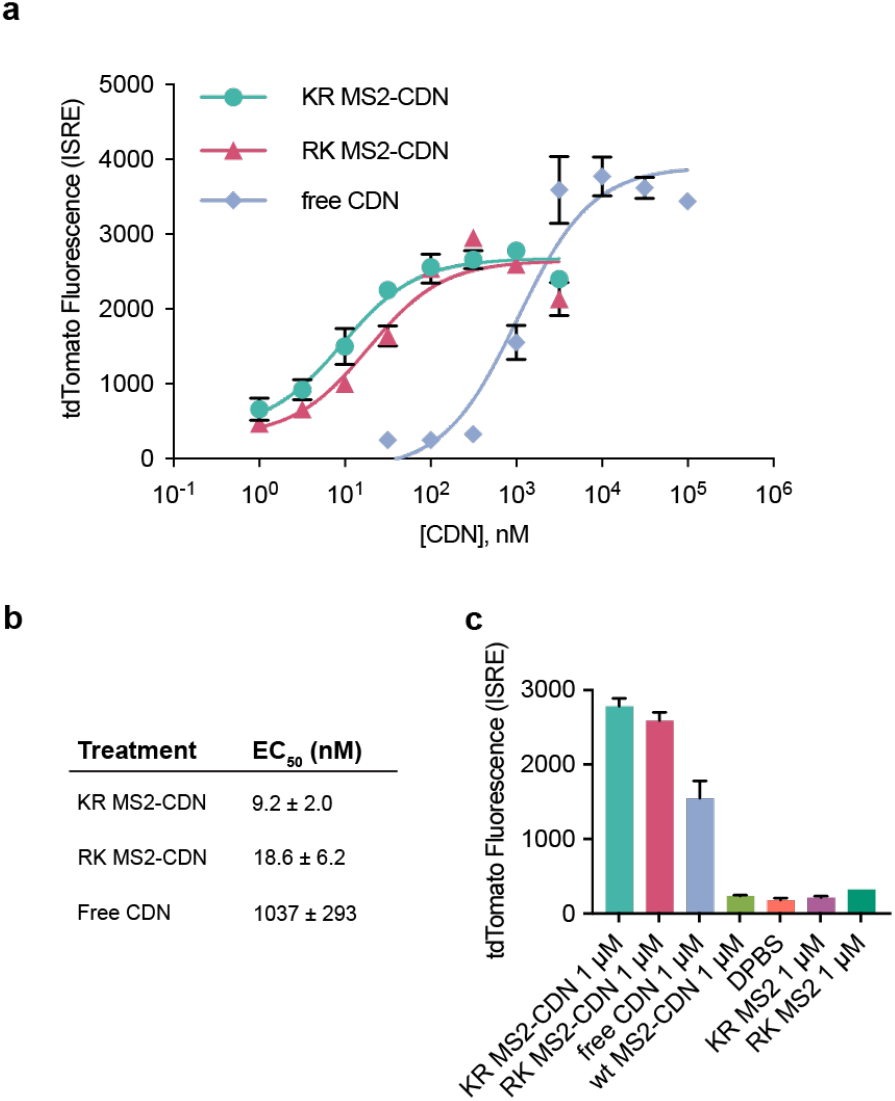
MS2-CDN treatment in a STING reporter line. **(a-b)** THP-1 STING reporter cells were treated with MS2-conjugated or free CDN in DPBS for 24 h. Red fluorescence was analyzed by flow cytometry and EC_50_ values were calculated from median fluorescence intensity values. **(c)** CDN formulations at 1 µM were compared to free MS2 and wild-type MS2 controls.

We observed that, after 24 h, the red fluorescence measurements at maximum treatment levels were consistently higher in cells treated with free CDN compared to those exposed to MS2-delivered CDN. We hypothesize that this was not caused by a reduction in the achievable degree of interferon production resulting from MS2 delivery. Rather, the previous naphthalimide fluorescence and mass spectrometric studies suggested a slower release of CDN from the capsid interior, which would result in a more gradual buildup of tdTomato. This could potentially be advantageous in providing a more gradual and sustained interferon response in target cells.

The low nanomolar EC_50_ values of MS2-delivered CDNs are a closer reflection of the natural binding affinity of the STING receptor to its CDN substrate, as well as the subsequent activation of STING ^44,45^. The 1 µM EC_50_ value of the free drug, on the other hand, more likely reflects the innate mechanisms limiting cellular uptake, such as the inability for CDNs to diffuse passively across the membrane and the relatively weaker affinity of CDNs to bind to anionic transport proteins on the cell surface. We hypothesize that the MS2 encapsulation and delivery of CDNs bypasses these negative regulatory elements.

One of the proteins known to aid in CDN membrane transport is SLC19A1^15,16^, which is known to carry folates, organic phosphates, and other anions into cells^46^. Using a SLC19A1 knockout version of the same THP-1 reporter cell line, a significant shift (p = 0.0242) in STING activation EC_50_ value was observed for the free CDN, indicating at least some reliance on SLC19A1 for cell entry. No such shift was observed for MS2-delivered CDN activity (**Figure 5A**), suggesting that the locally supercharged capsids can deliver CDNs directly to the cytosol without relying on endogenous membrane transport proteins. As SLC19A1 and other membrane proteins are inconsistently expressed across cell types and can be downregulated through resistance mechanisms in tumor cells, bypassing these membrane proteins could provide a substantial advantage to locally supercharged MS2 delivery systems for CDN STING agonists.

**Figure 5.**
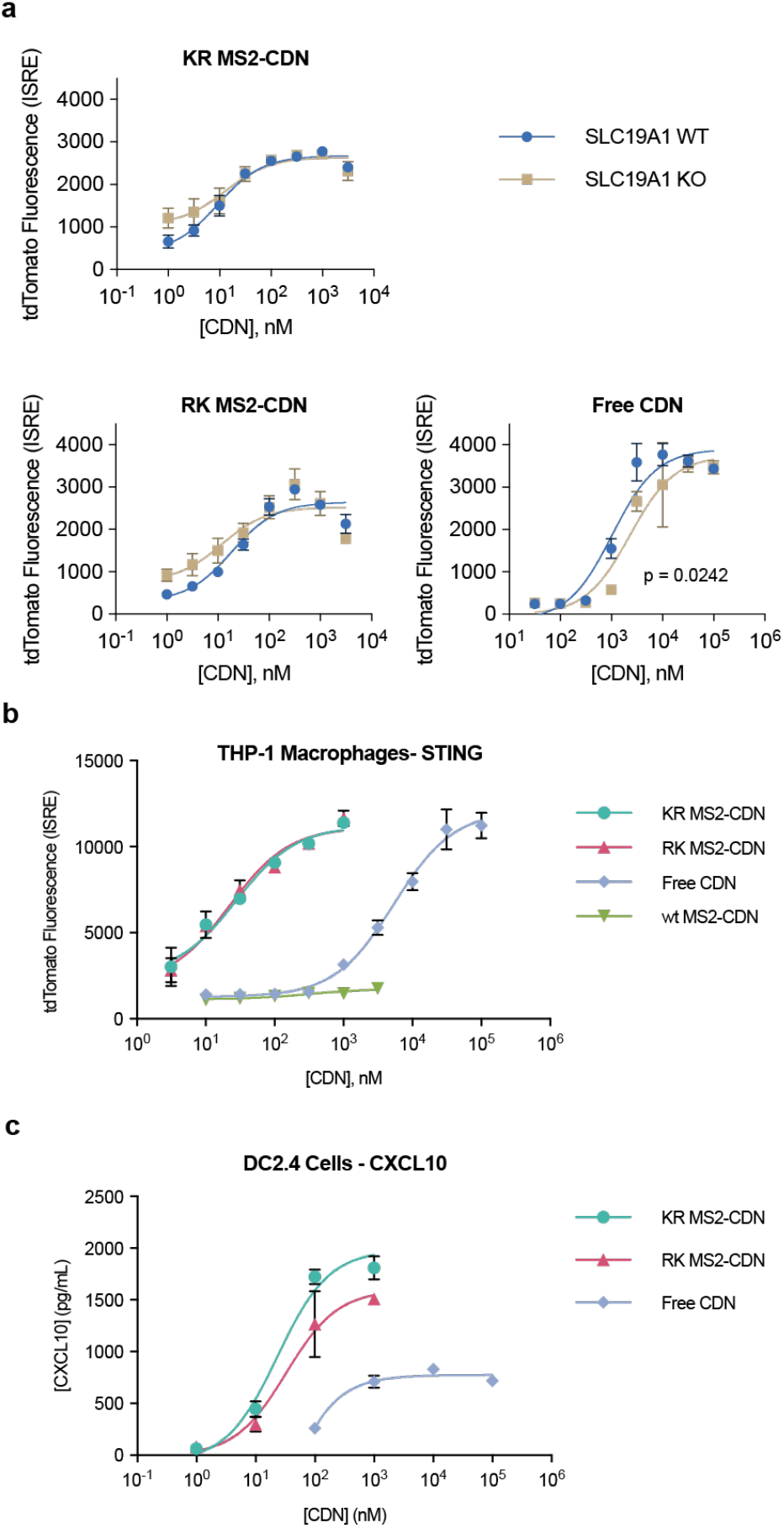
**(a)** STING activation after CDN treatment of THP-1 reporter cells with and without knockout of the SLC19A1 anion transporter. **(b)** STING activation after 24 h of CDN treatment of THP-1 reporter cells differentiated into macrophages by PMA treatment. **(c)** ELISA of secreted cytokine CXCL10 by DC2.4 murine dendritic cells after 24 h of treatment with free or MS2-conjugated CDN.

In addition to THP-1 monocytes, the MS2 delivery platform was tested on antigen-presenting cell types. The THP-1 reporter cells were differentiated into macrophages with phorobol 12-myristate-13-acetate (PMA) and again treated with free and MS2-delivered CDNs, showing a similarly improved STING activation profile for MS2-delivered CDNs as the undifferentiated monocytes (**Figure 5B**). To ensure that the CDN delivery was not caused simply by enhanced phagocytosis of the macrophage phenotype, MS2 capsid variants tagged with fluorescent dye were also delivered to both the monocyte and macrophage THP-1 cells, with their relative delivery profile remaining consistent (Figure S4). Enhanced downstream interferon response was also observed by STING activation in the mouse dendritic cell line DC2.4 (**Figure 5C**), which showed increased secretion of the cytokine CXCL10 at lower concentrations of CDN treatment.

## CONCLUSION

Cyclic dinucleotides and other small-molecule nucleotide and anionic drugs have considerable clinical potential in cancer therapies and other applications, but their development and systemic administration has been severely hindered by their poor or inconsistent uptake by cells of interest due to multiple endogenous or disease-specific regulatory pathways. Because of the low bioavailability of cyclic dinucleotide STING agonists, they must be administered at relatively high dosages, which can cause unwanted toxicity as well as increasing the risk of immune overactivation. This, in turn, could lead to adverse autoimmune responses—a persistent risk of some cancer immunotherapies^7,47^. By addressing the critical challenge of cell uptake, the locally supercharged MS2 drug delivery platform was demonstrated to be highly effective in delivering CDNs to cells with up to 100 times greater potency and downstream STING activation. Moreover, the use of covalent attachment to a monodisperse nanoscale carrier allows a defined number of drug molecules to be delivered through a specific mechanism-based release strategy. Taken together, this should lower the overall dosage level required to activate the STING pathway and achieve a downstream immune response, thus potentially expanding the therapeutic window for these compounds.

In terms of practicality, MS2 capsids are easily expressed and purified, and the sequence-defined uniformity of viral capsids allows for high batch-to-batch consistency. They have also been previously shown to exhibit minimal *in vivo* toxicity^48^. MS2 can be administered both as a monotherapy and in conjunction with other therapies, such as vaccines or checkpoint inhibitors that could be incorporated directly into the capsid scaffold through genetic fusion or previously developed bioconjugation strategies^41,49,50^. Further potential can be realized through the introduction of cell-specific targeting groups on the capsid surfaces, which is an area of ongoing research in our lab.

Finally, the MS2-based delivery platform can likely be adapted to other small-molecule nucleotide, oligonucleotide, or other anionic drug delivery applications. As anionic small-molecule drug candidates have been relatively underexplored in rational drug design efforts, an effective and generalizable cellular delivery system for anionic drugs could greatly expand their potential as effective therapeutic tools.

## ACKNOWLEDGMENT

This work was supported by the Panattoni Family as well as the Immunotherapeutics and Vaccine Research Initiative (IVRI). The authors would like to thank Professor David Raulet and his lab for sharing STING reporter cells and advice on STING agonist therapeutics, the UCSF Brain Tumor Center Preclinical Therapeutic Testing Core for sharing cancer cell lines, and Dr. Daniel Brauer for his advice and mentorship on MS2 expression and purification.

