## Supplementary Information for "Cytosolic delivery of anionic cyclic dinucleotide STING agonists with locally supercharged viral capsids"

#### Table of Contents

Supplementary Figures (S1-S4) …………………………………………………………………2
Materials and Methods ………………………………………………………………………… 6
Chemical Synthesis …………………………………………………………………………….. 9

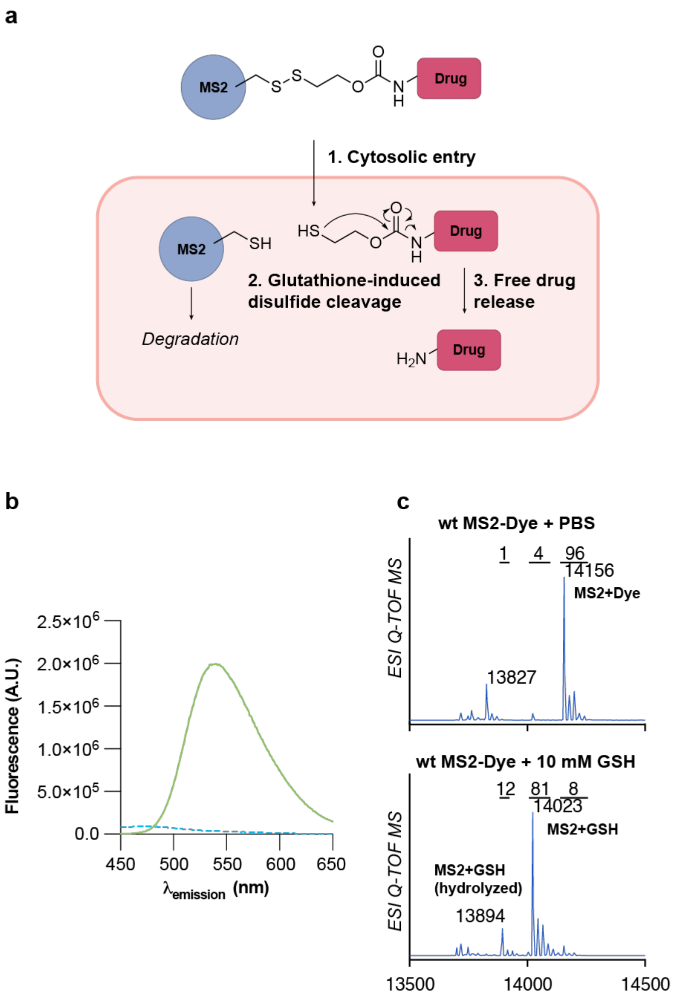


**Figure S1. (a)** Schematic of the disulfide-based drug delivery and release from MS2. **(b)** Fluorescence emission spectra of the naphthalimide dye in free and caged form. **(c)** Mass spectra of MS2-dye conjugate before and after treatment with 10 mM GSH.

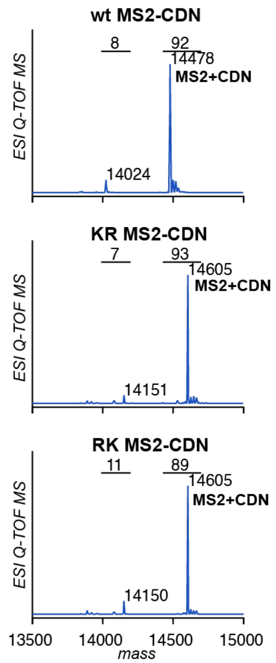


**Figure S2.** LC/MS-QTOF spectra of MS2-CDN conjugates.

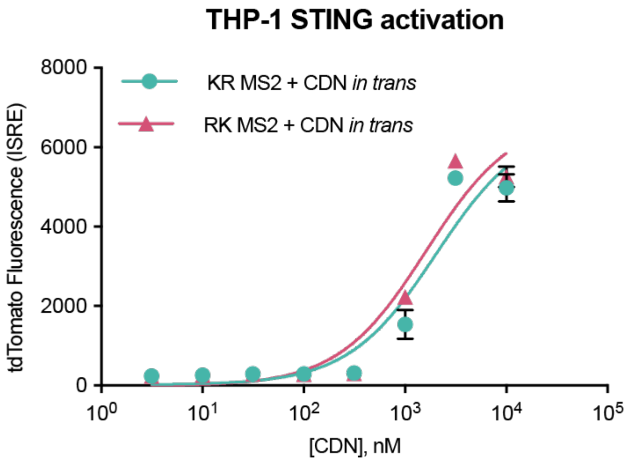


**Figure S3.** CDN drug delivery *in trans* of MS2. THP-1 STING reporter cells were treated with MS2 capsids and free CDN pre-mixed but not covalently conjugated for 24 h and analyzed for tdTomato fluorescence by flow cytometry.

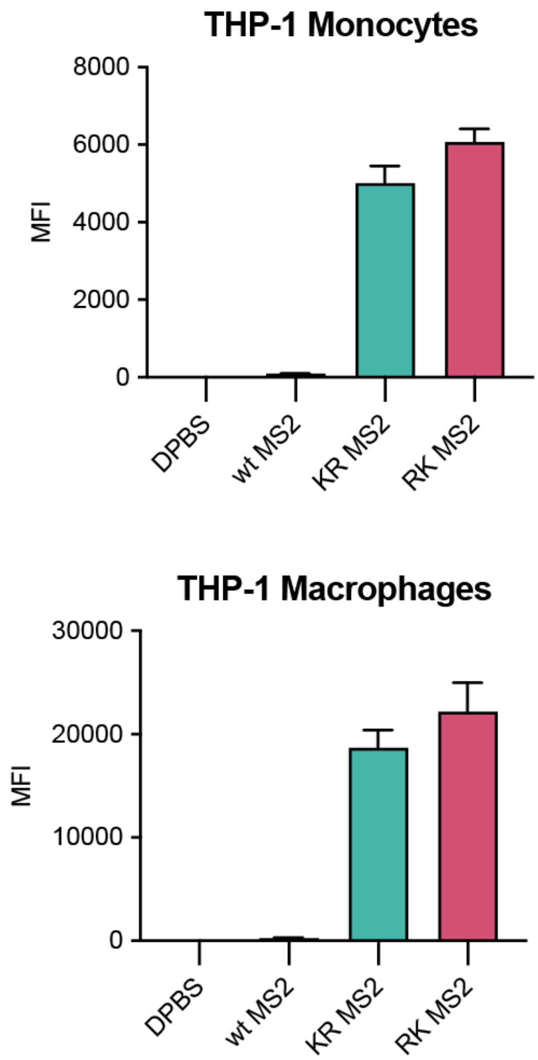


**Figure S4.** MS2 uptake in immune cell types. MS2 labeled with AlexaFluor 647 were administered to **(a)** THP-1 monocytes, and **(b)** THP-1 macrophages differentiated with PMA. All cell types were analyzed by flow cytometry.

**Materials and Methods:**

No unexpected or unusually high safety hazards were encountered. Cyclic dinucleotides were provided from Aduro Biotech. All other reagents were obtained from commercial sources and used without further purification unless otherwise indicated. Milli-Q $H_2O$ purified to a resistivity of 18.2 MΩ cm at 25 °C was used for all buffer and media preparations, and was obtained from a Barnstead NANOpure Diamond purification system.

**Equipment and instrumentation.** Protein purification was performed using a Cytiva AKTA go FPLC. Small molecule fluorescence spectra were obtained on a HORIBA Fluoromax-4 spectrofluorometer. Confocal microscopy was performed on a Zeiss LSM880 FCS at the Biological Imaging Facility (BIF), UC Berkeley. Flow cytometry measurements were performed on a ThermoFisher Attune NxT flow cytometer at the QB3 Cell and Tissue Analysis Facility (CTAF), UC Berkeley. Colorimetric measurements for MTS and ELISA assays were analyzed on a Tecan Infinite 200 PRO plate reader.

**Mass Spectrometry.** Purified proteins and small molecules were analyzed by electrospray ionization time-of-flight mass spectrometry (ESI-TOF-MS), which consisted of an Agilent 1260 series liquid chromatography (LC) system connected in line to an Agilent 6530 Q-TOF MS system. Samples were separated through a Proswift RP-4H monolithic analytical column and eluted using a mobile phase of nanopure $H_2O$ with 0.1% (v/v) MS grade formic acid (solvent A) and Optima MS grade MeCN with 0.1% (v/v) formic acid (solvent B).

**Expression and Purification of MS2 viral capsids.** MS2 capsid sequences were provided in a previous work and were purified as described in that work[1]. A 10 mL culture of each MS2 variant in DH10b *E. coli* was grown overnight in LB at 37 °C, then subcultured by a 1:200 dilution into 1 L of 2XYT media with 20 μg/mL chloramphenicol and grown at 37 °C until $OD_{600}$ reached 0.6, at which point cells were induced with 0.1% (w/v) arabinose and grown overnight. Cells were harvested by centrifugation, resuspended in 20 mM sodium phosphate, pH 7.4, and lysed by sonication. The supernatant was then collected, and protein was precipitated by addition of an equal volume of saturated ammonium sulfate solution and shaking overnight at 4 °C. The precipitate was then isolated, resuspended in 20 mM sodium phosphate buffer, pH 7.4, with 0.02% sodium azide, and filtered before FPLC purification with a HiScreen CaptoCore 700 column with an isocratic flow of the same buffer. Samples were then buffer exchanged to 1x phosphate-buffered saline via a HiPrep 26/10 desalting column. Purified MS2 was confirmed by SDS-PAGE and ESI-QTOF-MS and stored at 4 °C.

**Modification of MS2 internal cysteine by maleimide or pyridyl disulfide small molecules.** Each MS2 variant was diluted to 25 µM and mixed with 5-10 equivalents of small molecule in 200 mM phosphate buffer, pH 7.2, and allowed to sit overnight at 4 °C. Samples were then washed through seven rounds of filtration through Amicon Ultra 0.5 mL 100 kDa MWCO filters to remove excess small molecule and transfer into Dulbecco's Phopshate-Buffered Saline. Modification was confirmed by ESI-QTOF-MS.

**Mammalian Cell Culture.** HeLa, U-87-MG, U-251-MG, and RAW264.7 cell lines were cultured in DMEM + GlutaMAX supplemented with 10% FBS. THP-1 and DC2.4 cells were cultured in RPMI + glutamine supplemented with 10% FBS. Cell cultures were maintained at 37 °C in a humidified environment and 5% $CO_2$. Cell lines were validated regularly using short-tandem repeat profiling.

**AF594-conjugated MS2 Cell Uptake Assay.** N87C MS2 capsids were conjugated to AlexaFluor 594 maleimide as previously described[1]. For the free dye control, 50 µM AF594 maleimide was incubated with 100 µM glutathione overnight in DPBS at 4 °C to block maleimides. U-251-MG cells were plated in a 24-well plate at 100,000 cells/well and allowed to grow overnight. Cells were then suspended in 450 µL of growth media and treated with 50 µL MS2 or free dye to a final concentration of 5 µM dye. After 1.5 h, media was aspirated, and cells were washed three times with 20 U/mL heparin in DPBS to remove surface-bound MS2. Cells were then lifted with 40 µL 0.25% trypsin-EDTA, diluted in FBS-containing growth media, and analyzed immediately by flow cytometry.

**Cell viability assay.** Maleimide-val-cit-pAB-MMAE, Maleimide-val-cit-pAB-MMAF, free MMAE, and free MMAF were purchased from MedChemExpress. MS2-drug conjugates and free drug samples were filtered through a sterile 0.2 µm filter, and serial dilutions in PBS were prepared at 5x the desired treatment concentration. U-251-MG cells were plated in a 96-well plate at 4,000 cells/well and allowed to grow overnight. Media was aspirated and replaced with 80 µL media, and cells were treated with 20 µL of each drug formulation and incubated for 72 h at 37 °C. Media were then aspirated and the cells were resuspended in 200 µL MTS Assay Kit (Abcam, ab197010) diluted in DPBS and incubated for 30-60 min at 37 °C. The well plates were briefly shaken and analyzed by plate reader at 490 nm.

**Glutathione-triggered release assay.** MS2 containing disulfide-attached drug or dye was diluted to 10 μM in 50 mM phosphate buffer, pH 7.4. Glutathione was then added from a 100 mM stock solution to a final concentration of 10 mM, or an equal volume of water was added as a negative control. Dye samples were analyzed using a fluorometer with an excitation wavelength of 450 nm, and both dye and drug samples were analyzed by ESI-QTOF-MS. CDN drug samples were also analyzed by reversed-phase HPLC using $H_2O$ + 50 mM TEAA/MeCN as the mobile phase.

**Naphthalimide-conjugated MS2 Cell Uptake Assay.** HeLa cells were plated in 8-well chamber slides at 30,000 cells/mL and allowed to grow for 2 d at 37 °C. Cells were suspended in 180 μL DMEM without phenol red and treated with 20 μL of each dye-conjugated MS2 variant for 4 h. Cells were then imaged live by confocal microscopy, and images were processed via Fiji.

**STING agonist treatment of THP-1 cells.** THP-1 cells with a tdTomato STING reporter and (in some cases) a SLC19A1 knockout were provided as a gift from the lab of David Raulet, and had been generated in a previous study[2]. THP-1 cells were plated in a 48-well plate at 50,000 cells/well and treated immediately with a 10x dilution of MS2-conjugate or free STING agonist in DPBS. Cells were incubated at 37 °C for 24 h or analyzed over the course of 48 h for the time-course study, after which they were transferred to V-bottom 96-well plates and analyzed by flow cytometry. Data were analyzed with FlowJo software. Cells were gated by FSC/SSC, then the presence of a constitutive GFP signal indicating the presence of the STING reporter, before analysis of the tdTomato signal.

For STING analysis on THP-1 cells differentiated into macrophages, the same cells were treated with 125 ng/mL of phorobol 12-myristate 13-acetate and plated into a 96-well plate at 50,000 cells/well and differentiated for 48 h, after which differentiated cells adhered to the plate bottom. Buffer was then aspirated to remove dead or non-adhered cells, and fresh RPMI without PMA was added to recover cells for another 24 h, after which the differentiated THP-1 cells were treated with STING agonists and controls as previously. Prior to flow cytometry analysis, cells were lifted with 40 μL 0.25% trypsin and diluted with RPMI.

**Cytokine analysis of cell lines.** DC2.4 were plated in a 96-well plate at 50,000 cells/well and adhered overnight. Cells were then treated with a MS2-conjugate or free STING agonist in DPBS for 24 h at 37 °C. Cells were then centrifuged and the supernatants were collected. ELISA assays (Abcam) for CXCL10 and IFN beta were performed according to the manufacturer's instructions.

### Synthesis of linker-conjugated CDN:

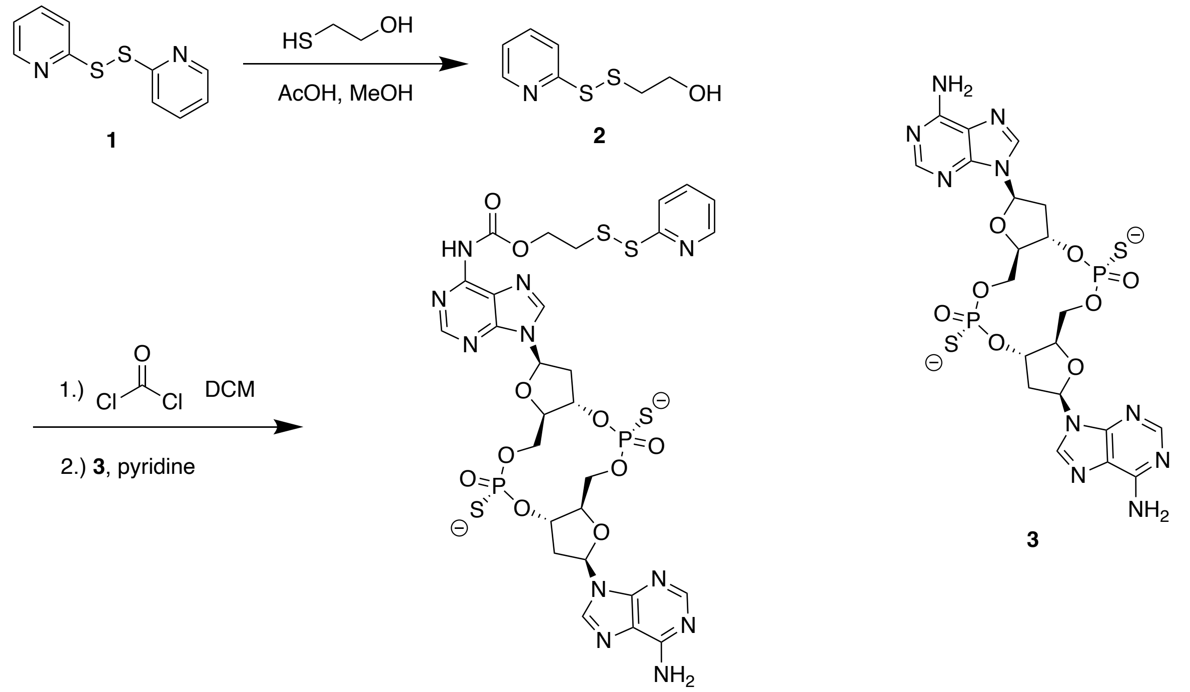


4

**Pyridyl disulfide 2**

A solution of 2,2-dipyridyl disulfide **1** (2.0 g, 9.08 mmol) was prepared in 25 mL of MeOH and 1.5 mL of AcOH. A 10 mL solution of 2-mercaptoethanol (320 µL, 4.53 mmol) in MeOH was then added dropwise over 10 min at room temperature, upon which the solution turned yellow. The reaction was stirred overnight at room temperature, after which the solvent was evaporated and the product was purified via flash chromatography (hexanes/EtOAc) to yield the product **2** as a clear oil (850 mg, 77% yield).

**CDN-disulfide conjugate 4**

Compound **2** (86 mg, 0.46 mmol) was dissolved in 1 mL of anhydrous DCM. A solution of phosgene (15% in toluene, 1.08 mL, 1.52 mmol) was added dropwise and the reaction was stirred at 0 °C under $N_2$ for 1 h. Solvent and excess phosgene were then removed by vacuum. About 1 mL of dry DCM was added and then removed under vacuum 3 times to remove excess phosgene. Cyclic dinucleotide **3** (10 mg, 0.015 mmol) was dissolved in 1 mL of dry pyridine and then added to the reaction mixture. The resulting reaction was stirred for 90 min at 0 °C under

$N_2$. The volatile components were then evaporated and the reaction was quenched with a 25% water solution in DMSO. The product was purified twice by Biotage C18 column chromatography (50 mM $NEt_3$•AcOH in $H_2O$/MeCN) with elution at 30% MeCN. Concentration of the fractions yielded **4** as a white solid (1.5 mg, 15% yield).

**Synthesis of linker-conjugated naphthalimide dye:**

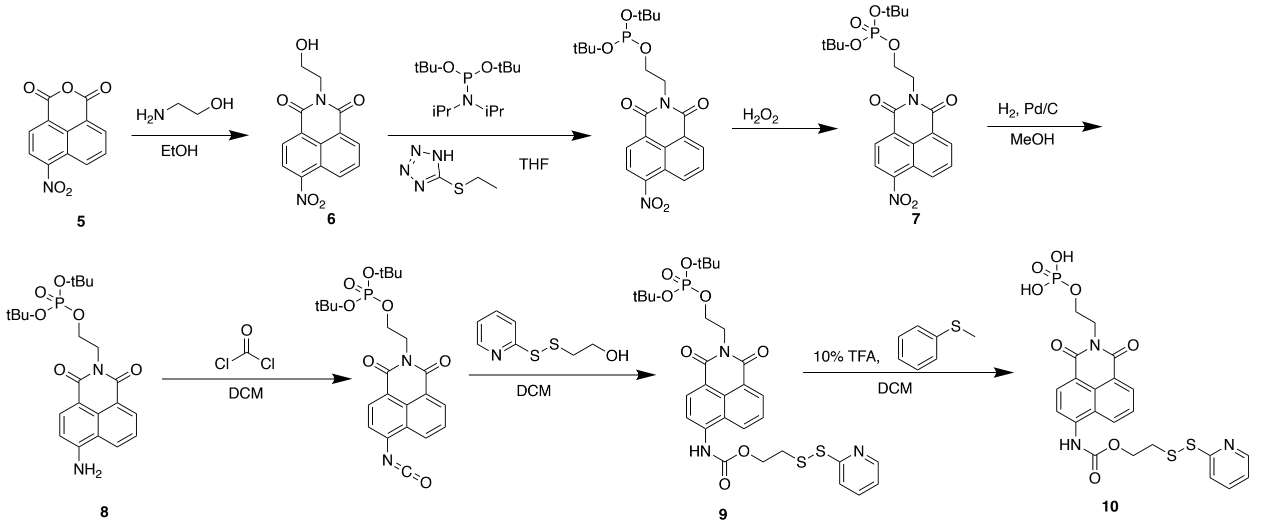


**Naphthalimide alcohol 6**

4-nitro-1,8-napthalic anhydride **5** (150 mg, 0.617 mmol) and ethanolamine (93 µL, 1.54 mmol) were combined in 5 mL of EtOH and stirred at 65 °C for 6 h. The solvent was then evaporated and the reaction mixture was dissolved in DCM and washed 3x with brine. The organic layer was then dried over $Na_2SO_4$ and concentrated. The product was isolated by flash chromatography (hexanes/EtOAc), yielding **6** an off-white solid (145 mg, 82% yield).

***Tert*-butyl protected phosphate naphthalimide 7**

Compound **6** (54 mg, 0.189 mmol), di-*tert*-butyl *N*,*N*-diisopropylphosphoramidite (238 µL, 0.755 mmol), and 5-ethylthio-1*H*-tetrazole (110 mg, 0.849 mmol) were dissolved in 1 mL of anhydrous THF and stirred overnight at room temperature under $N_2$. The reaction was then cooled to 0 °C and then 100 µL of hydrogen peroxide (34% in water) was added. The reaction was stirred for 3 h, then quenched with sat. aq. $Na_2SO_3$ and stirred for 30 min. The THF was evaporated, and the resulting mixture was extracted with EtOAc and washed with brine. The organic layer was dried over $Na_2SO_4$ and concentrated under reduced pressure. The product was purified by flash chromatography (hexanes/EtOAc) to yield **7** as a slightly yellow oil (35 mg, 39% yield).

**Amino naphthalimide 8**

Pd/C (5 mg) was wetted with EtOAc, after which 10 mL of MeOH was added slowly. A solution of compound **7** (35 mg, 0.073 mmol) in 10 mL of MeOH was then added. The flask was then evacuated and backfilled with $N_2$ three times before evacuating and bubbling in $H_2$ for 2 min. Over the course of the reaction the solution turned fluorescent green. The reaction was then stirred at room temperature under $H_2$ for 2 h, after which it was filtered through a bed of Celite to remove Pd/C, concentrated via rotary evaporation, and purified by flash chromatography (hexanes/EtOAc) to yield product **8** as a yellow-orange solid (34 mg, quantitative yield).

**Disulfide-conjugated *tert*-butyl phosphate naphthalimide 9**

Compound **8** (62 mg, 0.138 mmol) was dissolved in anhydrous DCM and stirred at 0 °C under $N_2$. Phosgene (15% in toluene, 690 μL, 0.968 mmol) was added dropwise, causing fluorescence to dim, and the reaction was stirred for 2 h. The volatiles were removed using a vacuum pump, after which 1 mL of dry DCM was added and evaporated 3 times to remove all excess phosgene. Alcohol **2** (182 mg, 0.969 mmol) dissolved in dry DCM was then added dropwise to the reaction while cooling to 0 °C. The resulting mixture was stirred overnight and warmed to room temperature under $N_2$. The reaction was then quenched with water, extracted with DCM, washed with brine, dried with $Na_2SO_4$, and concentrated under reduced pressure. The reaction mixture was purified via silica flash chromatography (hexanes/EtOAc) to yield **9** as a slightly yellow powder (34 mg, 41% yield).

**Disulfide-conjugated phosphate naphthalimide 10**

Compound **9** (24 mg, 0.0397 mmol) was dissolved in 10 mL DCM, and thioanisole (47 μL, 0.397 mmol) and TFA (30 μL, 0.397 mmol) were added sequentially at 0 °C. The reaction was stirred for 2 h, after which the mixture was concentrated and purified via C18 column chromatography to yield the phosphate-functionalized naphthalimide **10** as a slightly yellow solid (12 mg, 55% yield).
